# Experimental demonstration of daisy chain gene drive and modelling of daisy suppression systems

**DOI:** 10.1101/2025.09.20.677490

**Authors:** Jialiang Guo, Weizhe Chen, Jackson Champer

## Abstract

CRISPR-based gene drive can address ecological problems by biasing their inheritance coupled with an effector for either population modification of suppression. However, the risk of uncontrolled spread impedes some applications of gene drive. Daisy chain drives have received much attention as a potential approach to overcome this problem. They potentially allow efficient spread in a target population but are ultimately self-limiting. This is achieved by splitting a standard gene drive into multiple dependent elements, where each element can bias the inheritance of another, except one non-driving element. With the successive loss of each chain link, spread of transgenic elements will slow down and eventually stop. Here, we use modelling to assess the population dynamics of suppression daisy chain drives in both panmictic and continuous space models. We find that achieving population elimination through a single release of daisy chain gene drives is possible but difficult, with relatively high requirements for drive performance and release size. These effects are substantially amplified in spatial models. We also constructed two configurations of daisy chain gene drives in *Drosophila melanogaster* as a proof-of-principle. One is a rescue drive for population modification, and the other aims for population suppression by targeting a female fertility gene. Both functioned within expectations at moderate efficiency in individual crosses. However, the system failed to spread in cage populations because of higher than expected fitness costs. Overall, our study demonstrates that daisy chain systems may be promising candidates for both modification and suppression, but challenges remain in both construction and potential deployment in large regions.

## Introduction

Gene drives are selfish genetic elements that have the potential to spread through a population by being transmitted to progeny at super-Mendelian (>50%) frequencies^1–7^. They can be divided into modification drives and suppression drives based on their function. Modification drive enables a specific cargo gene to spread in the target population, thereby modifying the population for a particular purpose^2–4,6^, while suppression drive can directly eliminate the population, often by distorting the sex-ratio^8–11^, inducing sterility^12–15^, or both^1^^5^. CRISPR-based gene drives usually use Cas9 expression in the germline and gRNA(s) to induce DNA cleavage at the target site. The CRISPR homing gene drive will then be copied into the cleaved site through homology-directed repair, thereby converting heterozygotes for the drive allele into homozygotes in their germline. This allows biased inheritance. Such systems have been demonstrated in a wide range of organisms including yeast^16–19^, flies^14,20–25^, mosquitoes^26–28^, and mice^29^. These constructs have provided new approaches for reducing the transmission of vector-borne diseases, removing invasive species, and addressing agricultural pests^2–4,6,30^.

In CRISPR-based homing gene drive, the biased inheritance of drive elements relies on homology-directed repair. However, the cell has other repair options upon breakage. Non-homologous end-joining and microhomology-mediated end-joining pathways can potentially lead to sequence change, forming resistant alleles that cannot be targeted by drive elements. These represent a major challenge for homing drives. Resistance alleles are further categorized into two types. Nonfunctional resistance disrupts the function of the target gene, often imposing high fitness cost. On the other hand, functional resistance alleles will usually have higher fitness than a drive element^23^. Although this type of resistance allele is less common, it will outcompete the drive allele and cause drive failure^7,24,26,31,32^. One potential solution to prevent the formation of functional resistance alleles is to target a highly conserved region of an essential gene where mutation cannot be tolerated. By targeting such a gene while providing a recoded rescue to restore its function, nonfunctional resistance alleles can be purged because individuals inheriting them are often nonviable, and the risk of functional resistance (r1) can be further reduced by multiplexing gRNAs^3,7,13,33,34^. Functional resistance can also be addressed with gRNA multiplexing^14,24,31^.

If resistance can be addressed, CRISPR-based homing drives offer the potential to spread in a target population by a small release of drive individuals, thereby addressing some ecological issues efficiently. However, they also pose the risk of uncontrolled spread, leading to effects on non-target populations and even the risk of widespread species extinction^35,36^. Therefore, for several scenarios, developing temporally or spatially restricted gene drive systems could be a significant advancement in moving gene drives closer to implementation. Confinable gene drive systems can be spatially restricted. They will not spread in the population unless introduced above a frequency threshold. For example, CRISPR toxin-antidote gene drives use Cas9 and gRNAs as a “toxin”, which are programmed to cleave and disrupt an essential gene without copying the drive. The drive has a functioning copy of the target gene as an “antidote”, which is not be targeted by the gRNAs. Individuals who inherit disrupted alleles will be nonviable, but individuals with a drive allele will survive. Toxin-antidote gene drives have been tested successfully for modifying laboratory cage populations of *Drosophila melanogaster*^37–41^. Confined drive systems offer a more controllable option for gene drives, but they are still designed to persist in a population.

Another type of gene drive system that can avoid the risk of uncontrolled spread is self-limiting drive. Unlike introduction-threshold-based systems, which can persist when released above their threshold, self-limiting drives will eventually decrease in frequency, often after an initial increase, meaning that they will eventually disappear from the population over time^25,42–46^. However, such self-limiting drive systems often rely on repeated releases to have substantial suppressive effect, much like existing sterile insect technique (SIT) and release of insects carrying dominant lethals (RIDL) methods, with a single release usually failing to provide adequate power for successful suppression^47^. This can be a significant disadvantage when available resources are limited.

Daisy chain gene drive is a type of homing drive. It consists of sequentially linked drive elements. Each element can bias the inheritance of the next element, but the first element cannot have biased inheritance^44^. Eventually, the non-driving element will gradually be lost from the population, followed by sequential loss of other elements as they lose the ability for biased inheritance. This design is thus expected to initially spread efficiently, but eventually result in drive loss^44,48^. Since the concept of daisy chains was proposed in 2016, some studies have modelled the performance of daisy chains under different situations^44,48,49^, as well as the similar daisy quorum system^50^. The formation of resistance alleles can be a larger problem in daisy chains because increased cutting events can provide more opportunities for their generation^48^. Additionally, multiple DNA breaks induced at the same time can potentially lead to additional cellular stress and chromosomal abnormalities^5,51–55^, potentially inducing fitness costs that may have a greater effect on daisy chains than other homing systems. However, the handful of daisy drive studies have not assessed systems configured for suppression, nor have there been reports of examples in organisms.

In this study, we modeled the performance of suppression daisy chains in panmictic and spatial populations. We found that this kind of daisy chain system is highly susceptible to efficiency loss from both drive conversion rate and relative release size, but is less influenced by the embryo resistance rate (resistance alleles formed by maternal deposition of Cas9 and gRNA). We also constructed two daisy chains in *Drosophila melanogaster*, one of which targeted the haplolethal gene *RpL35A* with its last element while providing a recoded gene, aiming at population modification^34^. The other targeted the female fertility gene *stl* representing a population suppression drive^56^. Our daisy chains were able to induce super-Mendelian inheritance of all drive elements. However, even the more efficient modification drive failed to spread in a cage population. This can be attributed to the high fitness cost of the drive system, underscoring the need for daisy elements to be high-performance and fully characterized together.

## Methods

### Panmictic model

The performance of our three-element daisy chain drive was first tested in a panmictic population with non-overlapping generations through SLiM software (version 4.0.1)^57^. Our panmictic model is similar to previous studies^31,58,59^ and begins with reproduction, in which every fertile female will sample a male from the whole population. After the male is sampled, that chance that it is selected as the mate is equal to its fitness, which is decided by its genotype. If it is not selected, the female will attempt to sample another male. Each female has 10 opportunities to sample males, and if no male is selected, she will not reproduce. Fecundity in this model is based on both the population density and the fitness of this female, represented as:

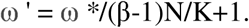

In this equation, N refers to the population size in this generation, β is the low-density growth rate in this population, which is set to 6 in our model, K is the population carrying capacity, and ω is the fitness of the female. The actual number of offspring produced follows a binomial distribution with n = 50 samples and a probability creating an offspring with each sample equal to p = ω ’/25.

In our model, three daisy element alleles are on three separate loci. We simulate them on either three independent chromosomes or far apart on the same chromosome by setting a recombination rate of 0.5 between each daisy element’s locus. In order to calculate genotype fitness for each individual, we assumed that each drive allele has an independent fitness cost (Table 1). The fitness of each individual is equal to the product of the genotype fitness of the three loci. For most alleles, we specify the fitness of homozygotes for the element (set to 0.98 by default, representing a small but present fitness cost). For heterozygotes, the element’s fitness is equal to the square root of the homozygote fitness. However, for a suppression element, fitness works differently. Female homozygotes are sterile, while heterozygotes have a separate fitness value, but only if a Cas9 element is also present. Male fitness is unaffected by suppression elements.

In our simulation, the non-drive element, Cas9 element, and effector element are modeled on three separate chromosomes. In germline cells, if an individual is a Cas9/wild-type heterozygote and has a non-drive element, then the wild-type allele will be converted into Cas9 for progeny according to the drive conversion rate. If an individual carries Cas9 and is an effector element/wild-type heterozygote, then drive conversion will similarly convert wild-type into effector alleles in the germline, which are then inherited by progeny. Both these processes can occur in the same individual for all three alleles are present, together with appropriate wild-type target alleles. If a wild-type allele is eligible for possible drive conversion but fails, it has a 50% chance to be converted into a resistance allele. Resistance may also form at the embryo stage at either potential site for any individual if the mother has Cas9 and a suitable gRNA for cleaving a particular wild-type allele (due to maternally deposited Cas9 and gRNA). The rate of this resistance allele formation is based on the parameter Embryo cut rate.

At the beginning of our simulation, we first allow a wild-type population consisting of 100,000 individuals to equilibrate for 10 generations. After that, we release a certain number of drive carriers which are homozygous for the non-driving element and the Cas9 element, but heterozygous for the last element. Released drive individuals have an equal proportion of males and females. In this study, we focus on three parameters that are most likely to affect the performance of daisy chains: drive conversion rate, embryo resistance rate, and female heterozygote fitness (in our study, this is a fitness cost affecting only Cas9-carrying females)^48^. In each simulation, we vary the release size (the proportion of released individuals to the total population) and one of the three parameters, while the other two parameters are kept at default values (Table 1).

**Table 1.** Default model parameters.

| Parameters | Default Values |
| --- | --- |
| Population carrying capacity | 100,000 |
| Drive conversion rate | 0.90 |
| Embryo cut rate | 0.10 |
| Female heterozygote fitness | 0.90 |
| Fitness (non-drive element) | 0.98 |
| Fitness (Cas9 element) | 0.98 |

### Genetic load

Genetic load in the field of gene drive is a characteristic that describes suppressive power on a target population. It measures the reduction in reproductive capacity in the population compared to a wild-type population of the same size, thus showing the effect of the drive^60^. Genetic load in this model is defined as:

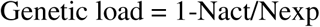

where Nact is the actual number of individuals observed in the next generation and Nexp is the expected number if all individuals were wild-type. To reliably measure the genetic load, we need to prevent population elimination from occurring during simulation (thus making an artificially robust population while still allowing accurate measurement of the genetic load). This is achieved by increasing the maximum offspring from 50 to 200 and elevating the low-density growth rate from 6 to 50. Due to the self-limiting nature of daisy chains, the suppression drive element can never reach a long-term equilibrium frequency. Therefore, we collect the maximum genetic load achieved in each simulation, rather than the more commonly reported equilibrium genetic load.

### Cage Simulation

SLiM was also used to model how our modification daisy chain will spread in a cage population. The simulation is based on our panmictic model with some changes on parameters. First, instead of a suppression element, the last element is from our haplolethal homing drive, which causes no sterility in females, but all individuals with a nonfunctional resistance allele are nonviable^34^. The population carrying capacity is set to 1,221, which is the average population during our cage experiment. In the first generation, the population size starts at 500, with 225 homozygote drive carriers. The drive conversion rate of the Cas9 element is set to 0.49, and the drive conversion rate of the modification element is set to 0.71 based on our experimental data. The germline resistance allele formation rate at the Cas9 locus was estimated by subtracting the drive conversion rate observed in our study from the published germline cut rate of Cas9 integrated at the same locus^34^. The embryo cut rate at Cas9 target site is set to 0.63 based on a previous study with a similar Cas9 element into *hairy* gene^61^. This site was also prone to functional resistance, so we assume that 1/3 of resistance alleles are functional (requiring a frameshift to prevent function). The embryo cut rate of the modification element was previously reported as 0.31^34^. However, that study observed a higher drive conversion rate than what we measured here, which indicates higher Cas9 activity. We therefore proportionally reduced the embryo cut rate. The germline resistance formation rate was similarly reduced. While functional resistance is possible from the recoded site at this gene, it is likely small^34,62^, and we set it to zero.

Because our initial model did not match our experimental cage data, we applied another parameter called homozygote transgene fitness. If an individual is homozygous for one of the three transgenes, we multiply its fitness value by this level (thus, a triple homozygote would have its fitness multiplied by the cube of this value). Heterozygotes at a site use the square root of this value (Table S1).

### Spatial model

We further extended our panmictic framework into a one-dimensional spatial model with a total length of 1. Unlike the panmictic model, in the spatial model, females can only sample mates from a with a restricted range with width equal to the *migration value*. Local competition between individuals will affect the fertility of females, rather than total population size. In order to quantify the influence of competition on female fertility, we defined a parameter called “expected adult competition”, which represents the expected density of individuals within a density interaction distance when the total population size is equal to the carrying capacity (*K*). This parameter is:

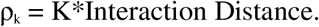

We also defined another variant ρ_i_ to measure the competition level within the interaction distance of each individual (ρ_i_). This is the local competition strength for each female within the interaction distance around her, which was set to 0.01. In this case, each individual contributes a strength from 1 to 0, which linearly decreases as its distance increases from 0 to 0.01. The fitness of a female is then defined as:

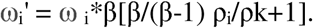

Offspring in our model are placed according to a normal distribution with a standard deviation scaled such that the mean displacement equals the *migration value.* If any offspring falls outside of the arena of length 1, we assign a new position until it falls within the boundaries.

We started the one-dimensional simulations by randomly scattering a population of K = 10,000 wild-type individuals into the whole space. After allowing the population equilibrate for 10 generations, we released drive individuals in the leftmost 20% of the area. The simulation was allowed to run for 50 generations

For an effective suppression drive, the population will undergo an initial decrease after drive release, followed by recovery as the drive declines in the population. To capture a suitable time scale for calculating the average population size, we recorded the number of generations it takes for the population size to recover to above 90% of the equilibrium size after experiencing a decrease (with the lowest population size less than 80% of the equilibrium population size). The longest recovery time among the three sets of parameters was ultimately used for our data output, which is 60 generations in our study. The maximum suppression range is measured using the suppression width. Specifically, in each generation, we divide the entire area into 50 slices. Then, we output the population size for each of the slices in every generation and identify slices where the population size is less than 10% of the average expected population size (indicating that 90% of the population is suppressed). We capture the generation which has the highest number of suppressed slices and find the total width of these slices.

### Plasmid construction

Our plasmids construction was partially based on EGDhg2t^38^, EGDh2^38^, and BHDaC^39^, which were constructed previously. The non-driving element was named as DN38C2 (daisy non-driving element inserted into the 38C region at *Drosophila melanogaster* chromosome 2 with two gRNAs). The Cas9 element was named as DHMDhCas9 (daisy homing modification drive inserted to *hairy* gene with Cas9 element). Reagents for restriction digest, PCR, and Gibson assembly, and plasmid miniprep were obtained from New England Biolabs and Vazyme; primers were from Integrated DNA Technologies Company BGI; 5-α competent *Escherichia coli* from Vazyme; and the ZymoPure Midiprep kit from Zymo Research. Plasmid construction was confirmed by Sanger sequencing. Plasmid sequences of the final drive insertion plasmid and target gene genomic region with annotation are provided on GitHub in ApE format^63^ (https://github.com/AlexGuojl/Daisy-chain-gene-drive-performance-modeling-and-demonstration-in-Drosophila-melanogaster).

### Generation of transgenic lines

Embryo injections were conducted by the UniHuaii Transgenic Flies Company. The first donor plasmid DN38C2 (300ng/ul) was injected into *w*^1118^ flies together with TTChsp70c9^64^ (300ng/ul) providing Cas9 and BHDacg1^39^ (300ng/ul) providing gRNA for transformation. The second donor plasmid DHMDhCas9 (300ng/ul) was also injected into *w^1118^* flies together with TTChsp70c9 (300ng/ul) providing Cas9 and EGDhg2t^38^ (300ng/ul) providing gRNA for transformation. Flies were housed in modified Cornell standard cornmeal medium (using 10 g agar instead of 8 g per liter, addition of 5 g soy flour, and without the phosphoric acid) in a 25C incubator on a 14/10-h day/night cycle at 60% humidity.

### Inheritance rate test

The drive conversion rate of the Cas9 element and final chain element (modification or suppression element) were tested separately. Initially, DN38C2 homozygous female virgin was crossed to DHMDhCas9 homozygous male (F0) to generate heterozygotes for both the non-driving element DN38C2 and Cas9 element DHMDhCas9 (F1). The same method was used to generate DHMDhCas9-HSDstlU4 and DHMDhCas9-AHDr352V2 heterozygotes. The F1 males or virgin females were collected, and then single-pair mated to *w^1118^* individuals of the opposite sex, and resulting F2 progeny were assessed for efficiency of drive elements. F1 parents were allowed to lay eggs for 7 days and then removed. F2 progeny were collected every two days from day 12 to day 18 after crossing.

Flies were anaesthetized with CO2 and screened for fluorescence using the NIGHTSEA adapter SFA-GR for DsRed and SFA-RB-GO for EGFP. A 3xP3:EGFP cassette was used to indicate the presence of the Cas9 for DHMDhCas9. 3xP3:dsRed cassettes are used to indicate the presence of HSDstlU4 or AHDr352V2. Cassettes driven by the 3xP3 promoter will lead to expression in eyes, which is easy to visualize in the white eyes of *w^11^*^18^ flies. A polyubiquitin promoter:dsRed cassette was used to indicate the presence of DN38C2. This cassette will lead to an expression of DsRed at the entire body.

### Phenotype data analysis

To estimate drive inheritance, drive conversion, and related parameters, we performed a pooled analysis (Data Sets S1-3).), combining data across all individual crosses. However, this approach does not account for potential batch effects, as each cross may represent a distinct batch with unique characteristics, potentially biasing rate and error estimates. To address this, we fitted a generalized linear mixed-effects model with a binomial distribution using maximum likelihood estimation (Adaptive Gauss-Hermite Quadrature, nAGQ = 25), allowing for variance between batches^31,34^. This approach yields parameter estimates that are generally consistent with the pooled analysis but with slightly higher standard errors. All analyses were conducted in R (v4.4.2). These results are featured in the manuscript.

### Cage study

Flies were housed in 25x25x25 cm enclosures with nine food bottles for the cage study. We initialized the cage population by adding 25 homozygous drive carrier females and 31 *w^1118^* females to each food bottle, which gives us 225 drive carrier females and 279 *w^1118^*females for each cage. Each female individual was previously allowed to mate with males of the same genotype for two days before being released into the bottle. Females were allowed to lay eggs in bottles for 24 hours. Then, we removed the females and unplugged the bottles. After 11 days, we remove all adults from the bottle and replace the old bottles with new bottles. All adults were allowed to lay eggs for 24 hours and were then removed and kept them for phenotyping. We repeated these steps every 12 days.

### Fertility and viability test

Triple heterozygote males were crossed one to one with *w^1118^* females. Female progeny that were either *w^1118^* (though possibly heterozygous for resistance alleles) or that were triple daisy element heterozygotes were collected. After three days, we conducted one-to-one crosses for those females with *w^1118^* males, which are also generated from one-to-one crosses between two *w^1118^* parents. Each pair were in different vials, and we counted the number of eggs after 24 hours. We then move each pair of parents into a new vial and again counted eggs after 24 hours. We repeated this three times for each pair of parents. After that, we phenotyped offspring from each vial from day 12 to day 18 after parents were removed.

## Results

### Performance of suppression daisy chain drives

We first modeled the performance of a suppression daisy chain drive in a panmictic population. In this design, all three elements are genetically unlinked (located at different chromosomes or far apart on the same chromosome). With the help of the Cas9 element, the gRNAs expressed by the non-drive element can bias the inheritance of Cas9 element, and the suppression element will bias the inheritance of itself when together with Cas9. Any female that carries two copies of the suppression element will be unable to produce offspring.

Nonfunctional resistance alleles at the Cas9 site are recessive lethal, and nonfunctional resistance alleles at the suppression target site are recessive female sterile. In a daisy suppression drive, the non-driving element will be more rapidly lost than in a modification system because it will often find itself in sterile females, followed by the Cas9 element once it no longer is able to frequently bias its own inheritance with the help of the non-drive element (Figure 1). Finally, the suppression element will decline. This is in contrast to a self-sustaining homing suppression drive, which will increase to an equilibrium frequency and remain there in the absence of functional resistance allele formation.

**Figure 1.**
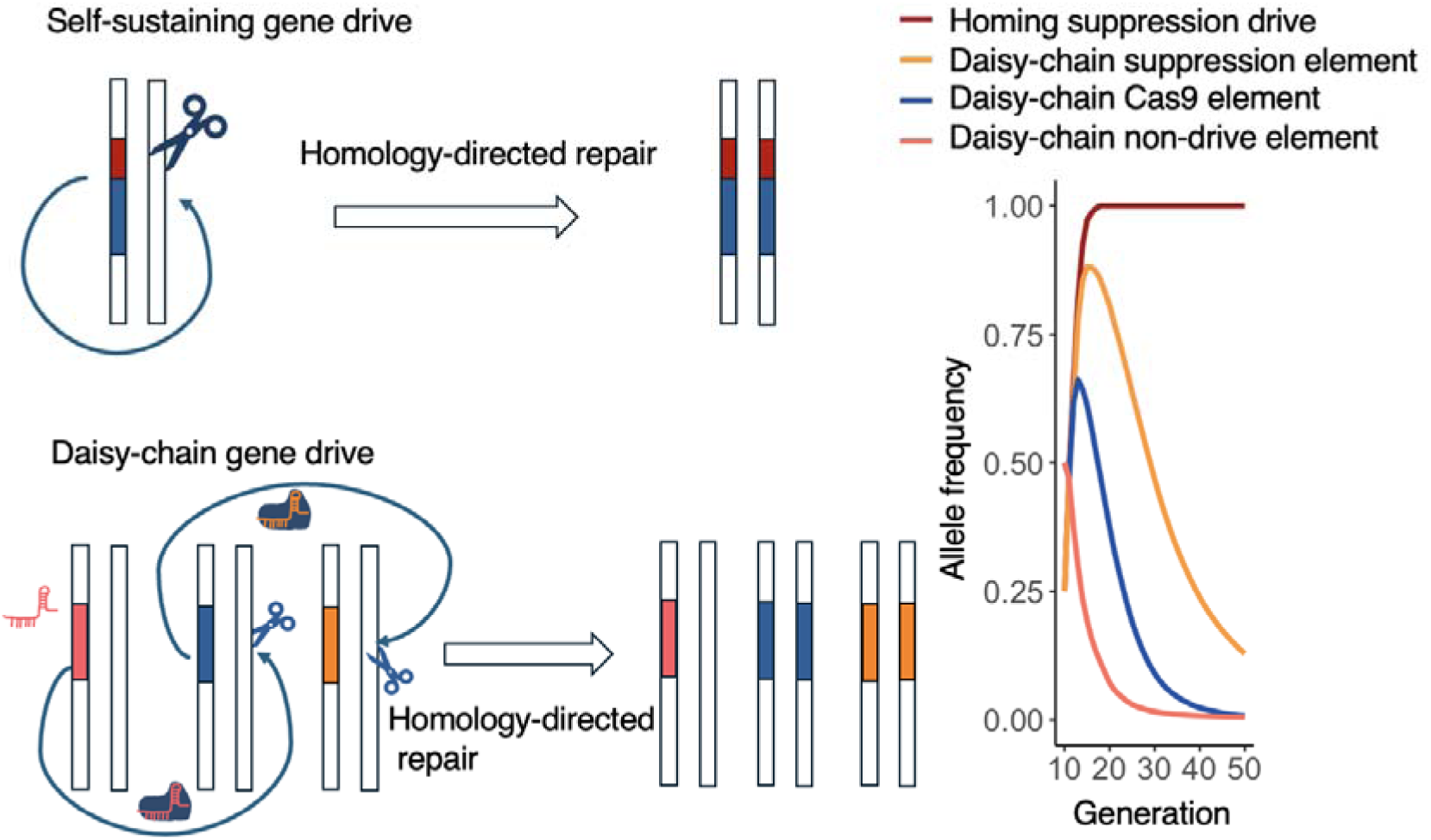
Mechanism of the daisy chain drive. In self-sustaining homing drive, the drive contains all necessary components to copy itself. In an optimal 3-element daisy chain drive, the gRNA expressed by a non-drive element targets the wild-type allele opposite the Cas9 element, and the gRNA expressed by the final element (for suppression in this case) targets the wild-type allele opposite itself. The line chart shows the change in allele frequency in a panmictic population with 100,000 wild-type individuals. With ideal drive performance, the initial population is composed of 50% wild-type and 50% of individuals that are heterozygous for the suppression element and homozygous for other elements. The cas9 and suppression element frequency increases at first, but eventually all elements disappear from the population, unlike the self-sustaining drive.

We modeled a high performance but imperfect daisy suppression drive system (Table 1). Our Panmictic simulation results shows that our daisy chains require large (single) release sizes to achieve success. When the release size is below 60%, the drive will never eliminate a population under any combination of parameters included in our simulation (Figure 2a-c).

There is also a threshold of conversion rate at 70% under which the drive cannot eliminate the population (Figure 2a). Interestingly, when the conversion rate was above this threshold, the success rates of population suppression decreased as the conversion rate increased in the case of intermediate release size. We tracked the allele frequency of both the Cas9 element and suppression element to explain this (Fig S1). Our result shows that a high conversion rate with a medium release size will lead to a faster reduction in population size, but also even more rapidly reduce our non-driving element and Cas9 element. Thus, in later stages, the suppression element lacks enough driving force to stay at high frequency and eliminate the population, despite reaching a higher peak frequency and maximum genetic load (suppressive power) (Figure 2d). On the other hand, a moderate conversion rate will be able to decrease the population size over a longer period of time, even though it’s maximum level may be slightly lower, thus eventually eliminating the entire population.

Unexpectedly, embryo resistance rate had only a small influence on the success of a suppression daisy chain in our simulations (Figure 2b). For self-sustaining suppression drives, embryo resistance can be highly detrimental^65^. Previous research on modelling modification daisy chains has shown that major changes in the maternal deposition/embryo resistance rate can have a significant effect but be compensated for with relatively minor changes in the release frequency^48^. Compared with modification daisy chains, suppression daisy chains can succeed only in a narrow range of relatively high release size. Therefore, it is possible that the impact of maternal deposition in suppression daisy chains is already compensated by the high release size required for achieving population suppression. Though resistance allele generation caused by maternal deposition can slow the drive, it can also rapidly make additional females sterile, even if they don’t have two drive alleles. The success or failure of a self-limiting suppression drive often occurs within a short period of time after the drive is released. Thus, the effects of embryo resistance may largely cancel out. Note that for genetic load, the detrimental effects of higher embryo resistance are more apparent at low release frequencies (Figure 2e)

As expected due to their direct negative effects, female fitness costs from the suppression element also significantly influences the success rate and maximum genetic load of suppression daisy chains (Figure 2c,f). As the release size decreases, the drive requires a lower fitness cost to achieve population elimination.

**Figure 2.**
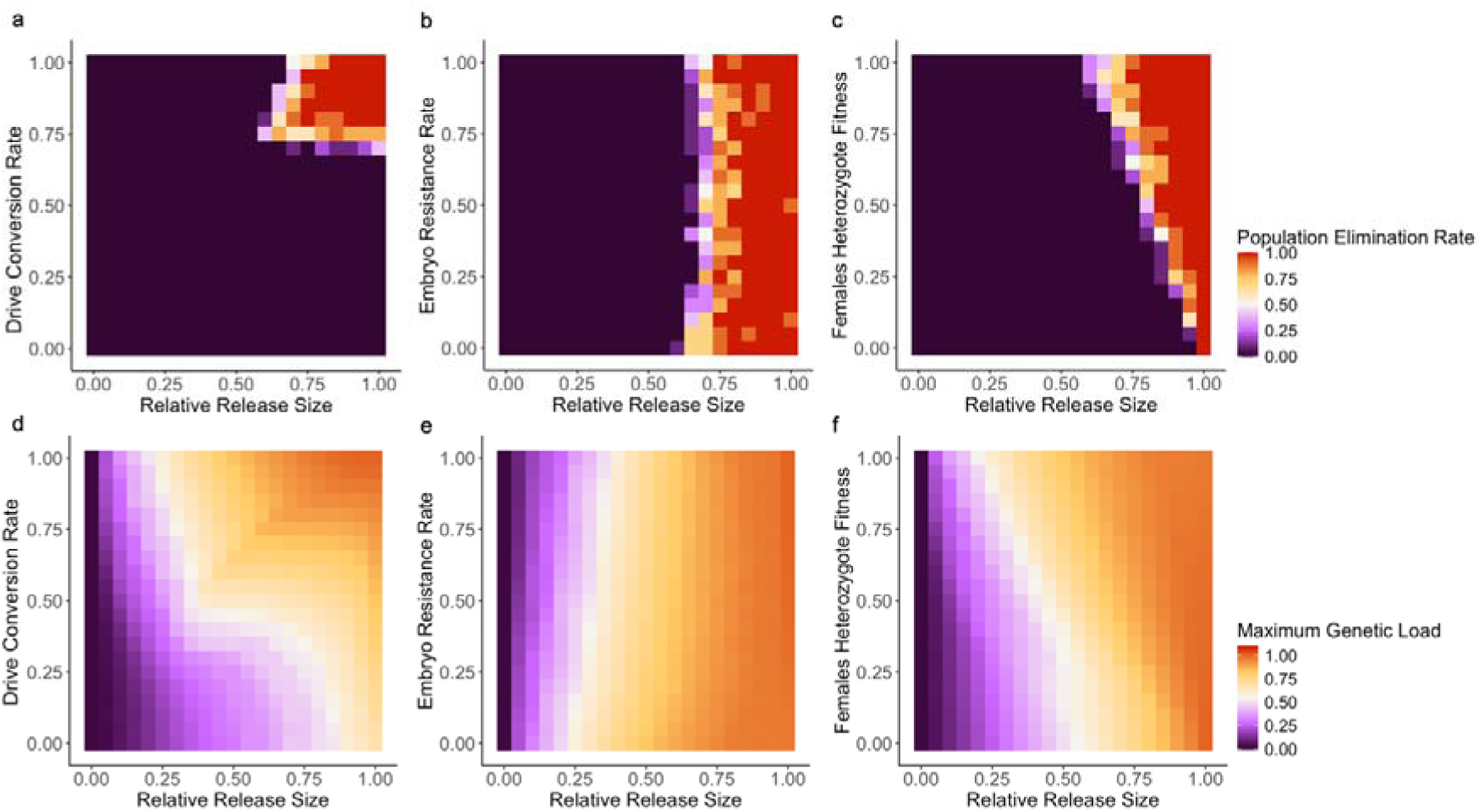
Performance of suppression daisy chain drive. For daisy suppression drives, we measured the a-c rate of successful population elimination within 50 generations and d-f maximum genetic load (suppressive power). We varied the release size (fraction of the population that initially contained the drive) and the a,d drive conversion rate, b,d embryo resistance rate, and c,e fitness of female heterozygotes. Each point shows the average of 10 replicates.

### Spatial performance

Because self-limiting suppression drives tend to decline rapidly in frequency, they may be unable to spread effectively in spatial environments. To investigate this, we uniformly released drive individuals in the left 20% of an arena. The width of suppression was defined as the region with at least a 90% decrease in population size. Our results indicate that the drive allele cannot significantly spread beyond the boundary of the release region, even when the release size is well above the threshold for panmictic population elimination (Fig 3).

Within the release region, a high conversion rate beyond 40-60% does not lead to a better result in terms of reducing the average population (Fig 3b). On contrary, a moderate conversion rate, which is unable to eliminate the population in panmictic simulations, can allow the suppression element to exist in population for a longer period and therefore facilitate longer term reduction in the population size (Fig S2). However, the peak population reduction tends to still be lower when drive conversion is high, both for total population reduction (Fig 3a) and width of high population reduction (Fig 3e).

As expected, lower embryo resistance (Fig 3c) and higher female heterozygote fitness (Fig 3d) results in significantly improved suppression due to the drive’s greater persistence, somewhat differing from the result of embryo resistance in the panmictic model. However, the maximum width of heavy suppression was not substantially affected (Fig 3f-g). This is because the width of suppression is an instantaneous value, which only includes the situation of a certain generation with the strongest suppression effect. High embryo resistance and low somatic fitness tends to induce strong but more transient suppression due to more rapid elimination of the drive despite additional temporary suppressive power from nonfunctional resistance alleles that result in female sterility or other fitness costs in drive females. Overall, even under the ideal parameter combinations, daisy chain drives can only lead to a modest and short-term decrease in population size, and even then, only in the area where they are released. Rapid elimination of transgenic elements coupled with dispersal of wild-type individuals prevents success from a single release, though sustained or widespread releases would be able to restore efficacy.

**Figure 3.**
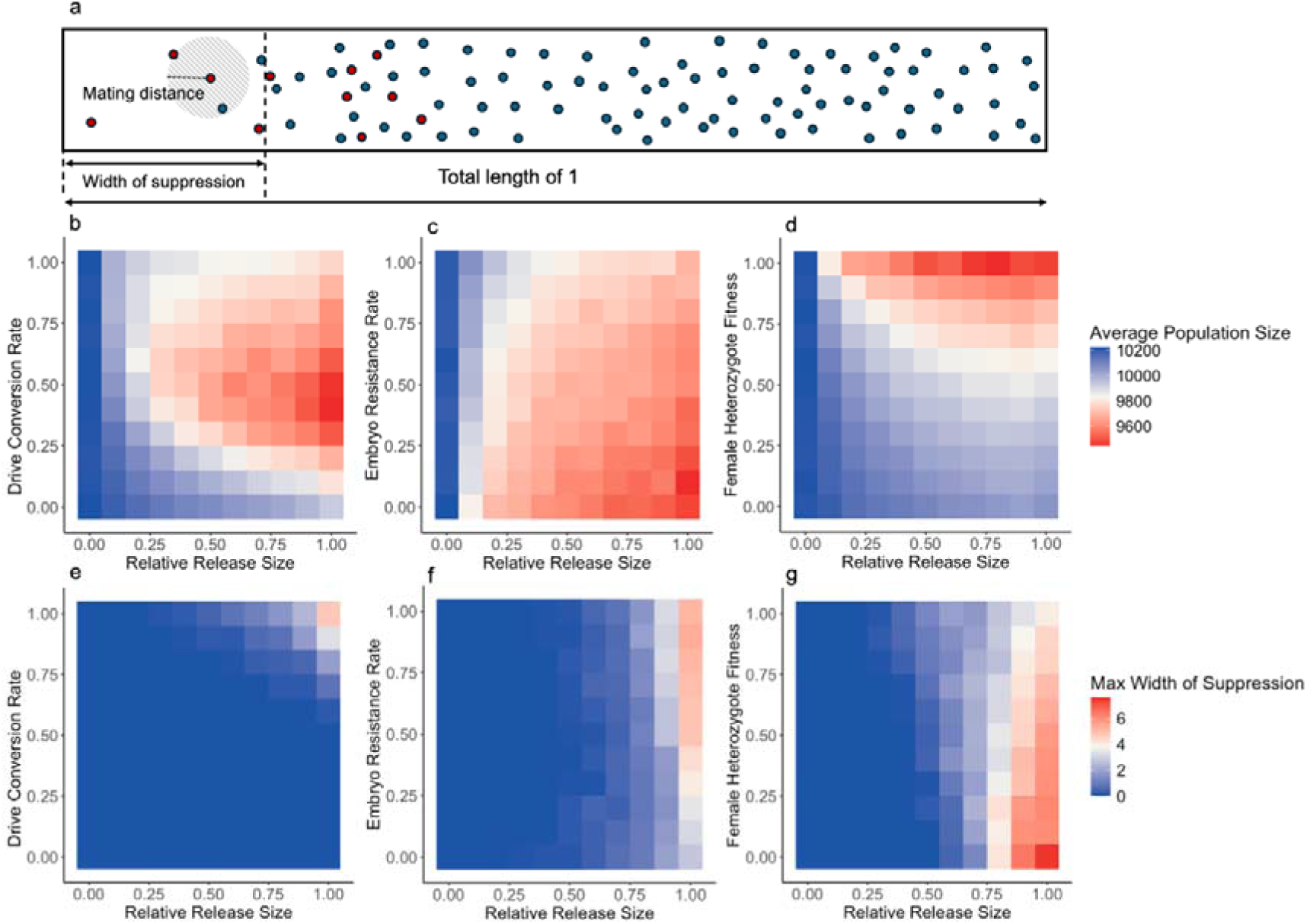
Suppression Daisy chains in spatial populations. **a** In one-dimensional space, drive carriers were released in the leftmost 20% of the arena (the relative release size shows what fraction of individuals in this area were initially drive). Red dots represent drive carriers and blue dots represent wild-type individuals. In this space, each female will mate and distribute offspring only with within a limited distance. We measured the **b-d** average population size of the whole arena and **e-g** greatest width with at least 90% population decrease in each 2% of arena width. We varied the release size (fraction of the population that initially contained the drive) and the **a,d** drive conversion rate, **b,d** embryo resistance rate, and **c,e** fitness of female heterozygotes. Each point shows the average of 10 replicates.

### Drive construct design

In this study, we designed two daisy chains for population modification and population suppression. Our suppression and modification drive constructs include three elements. Both designs use the same non-driving element and Cas9 element, differing only in the final element. The non-driving element was inserted into a genomic region located between two genes on chromosome 2L to minimize potential interference with native genes and therefore avoid affecting an individual’s fitness (this location was also used for drives in a previous study^31^). This element consists of two gRNAs to increase drive efficiency and reduce the functional resistance allele formation rate by providing two opportunities for cleavage. It has polyubiquitin-DsRed as a marker, producing red fluorescence in the body. The non-driving element targets the first exon of the *hairy* gene, a haplosufficient gene where null alleles are recessive lethal^38,66^. This biases the inheritance of Cas9 element, which is located at the gRNA’s target site. The Cas9 element contains a recoded version of the *hairy* sequence, allowing it to remain viable. Cas9 expression is controlled by the *nanos* promoter and UTRs, and an EGFP is expressed by a 3xP3 promoter, producing green fluorescent protein expression in eyes.

Our modification element and suppression element were constructed previously^34,56^. They are marked by a DsRed fluorescent marker driven by the 3xP3 promoter for expression in the eyes to indicate the presence of a drive allele. The modification element includes two gRNAs targeting exon 4 of a highly conserved haplolethal gene *RpL35A*, a protein component of the 60S ribosomal subunit^67^. This element also includes recoded “rescue” version of *RpL35A*. The suppression element targets *stall* (*stl*). Null homozygous mutations at this locus result in sterile females but show no effects on males. The suppression element contains four gRNAs within tRNA scaffolding that target the fourth exon of *stall*.

**Figure 4.**
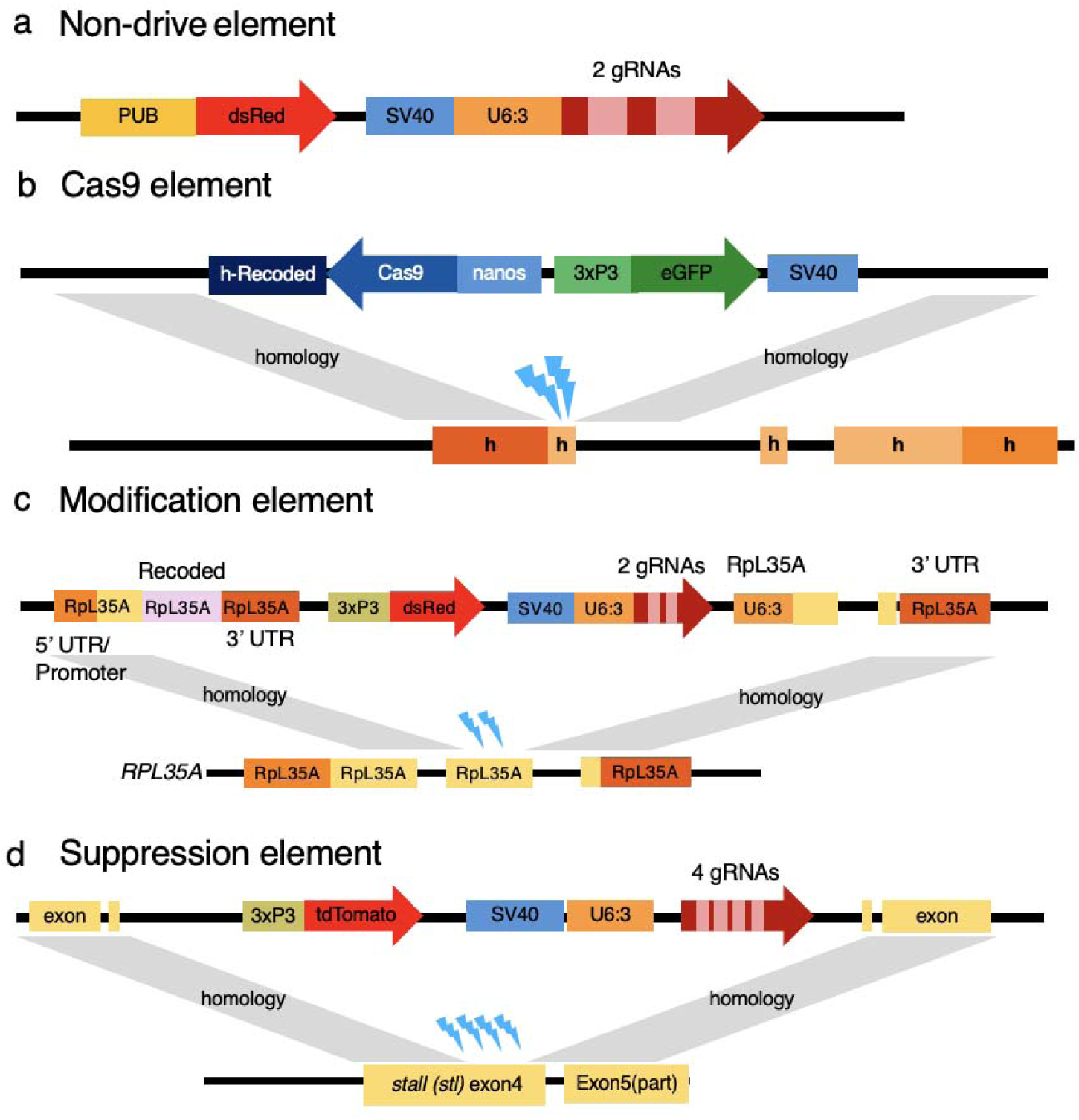
Construction of modification and suppression daisy chain drives. **a** The non-drive element contains a DsRed marker under the control of the PUB promoter for expression in the entire body. The two gRNAs target *hairy*, facilitating the homing of the **b** Cas9 element. Cas9 is driven by the *nanos* promoter, and this element also contains a recoded version of *hairy* and an eGFP marker. **c** In the modification daisy chain design, the Cas9 element will combine with two gRNAs in the “effector” element targeting the second coding exon of RpL35A. **d** In the suppression daisy chain design, the suppression element has four gRNAs targeting the fourth coding exon of *stall* gene. Both effector elements contain a red fluorescent marker driven by the 3xP3 promoter for strong expression in the eyes.

### Drive performance assessment

We first tested the inheritance of different drive elements separately by crossing double heterozygotes with *w^1118^* individuals. All drive individuals received all of their transgenes from a male parent. For individuals heterozygous for the non-drive and Cas9 elements, progeny of females showed a 72% Cas9 element inheritance rate, and the progeny of heterozygous males showed a 58% inheritance rate (Fig S3). For individuals heterozygous for the Cas9 and effector elements, the inheritance rate of the modification element was 83% among progeny of males and 85% among progeny of females. The inheritance rate of the suppression element was 89% among progeny of females and 82% among progeny of males. The inheritance rates of all the daisy chains elements were significantly higher than the 50% expected under Mendelian inheritance (*P* <0.0001, z-test, Data Set S1).

We then tested the inheritance rate of different elements in triple heterozygotes that received all their transgenes from male parents. Both modification daisy chains and suppression daisy chains induced biased inheritance of drive elements (*P* < 0.0001, z-test, Data Set S2) (Fig 5). For the modification and suppression elements, drive inheritance was over 80%. For the Cas9 element, inheritance rates in females reached nearly 80% but were lower than 70% in males. No biased inheritance was found in the non-drive element for either cross element.

**Figure 5.**
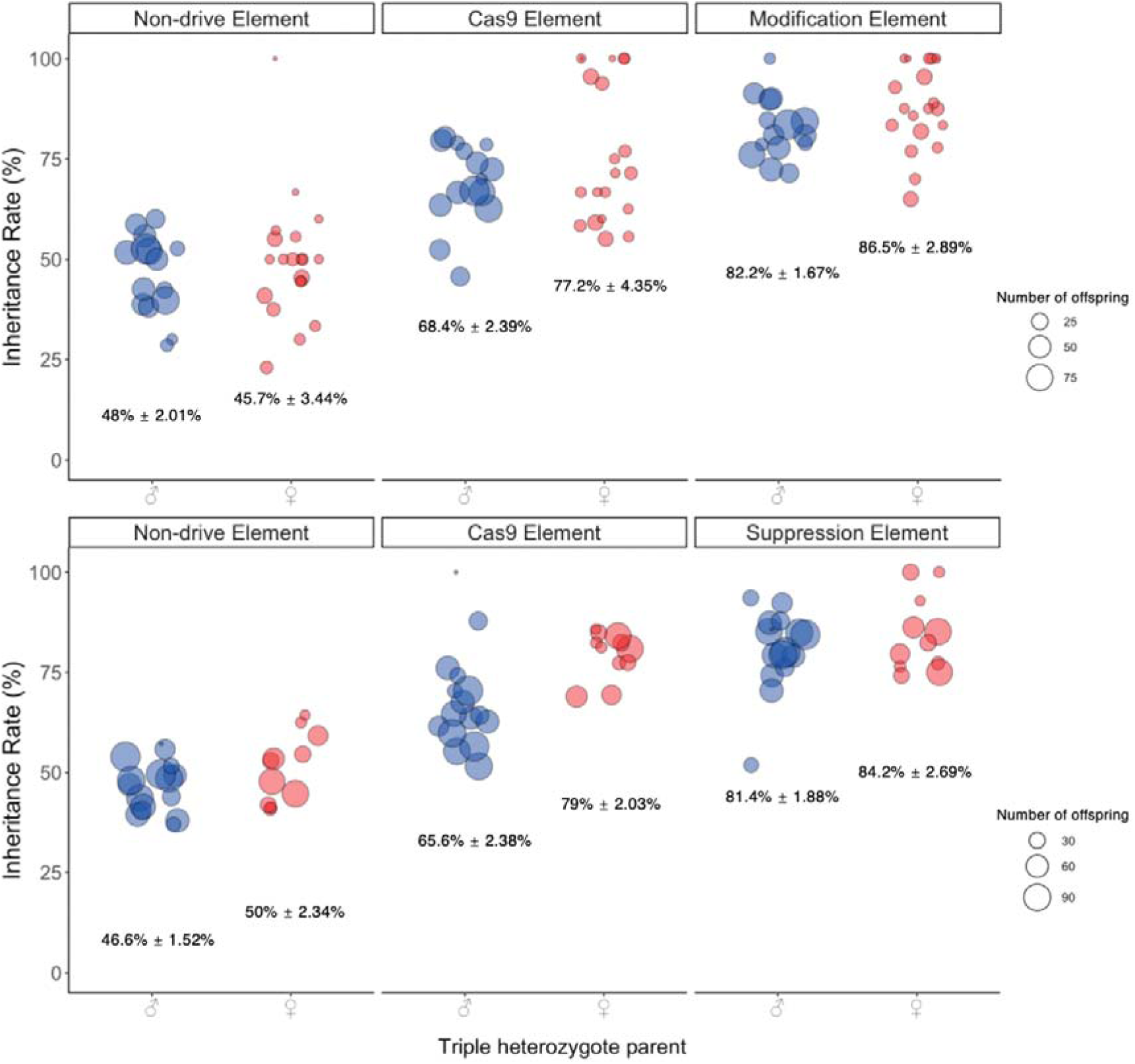
Daisy chain drive performance. Flies heterozygous for the non-drive, Cas9 and effector (either modification or suppression) elements (with all transgenic elements inherited from their father) were crossed with *w^11^*^18^ flies. The inheritance rate of each allele in offspring was evaluated based on their fluorescence. Each data point represents the inheritance rate observed in one vial containing one drive parent, and the size of each dot represents the clutch size representing the total number of adult offspring phenotype. Also shown are the mean and standard error.

Overall, this indicates that despite potential gRNA saturation effects, the daisy drives operated with good efficiency.

### Daisy drive cage study

To further explore the performance of daisy chain drives in a relatively large population, we performed a cage study. The modification drive was selected because the suppression system was not likely to have enough power to eliminate the population, even when assessed as a self-sustaining system^56^. Surprisingly, drive performance in the cages was poor, with all elements immediately decreasing in carrier frequency despite a homozygous release (Fig 6). After just four generations the Cas9 carrier frequency decreased from 44% to 13%, and the modification element frequency decreased from 44% to 21% (Data Set S3). Based on a SLiM model with parameters derived from experiments, which included a moderate fitness cost for Cas9, we expected the Cas9 element and especially the modification element to rapidly increase in frequency (Fig 6). The model was close to the cage data only when a heavy homozygous fitness cost of 78.9% was applied to each of the three elements of the daisy drive system.

**Figure 6.**
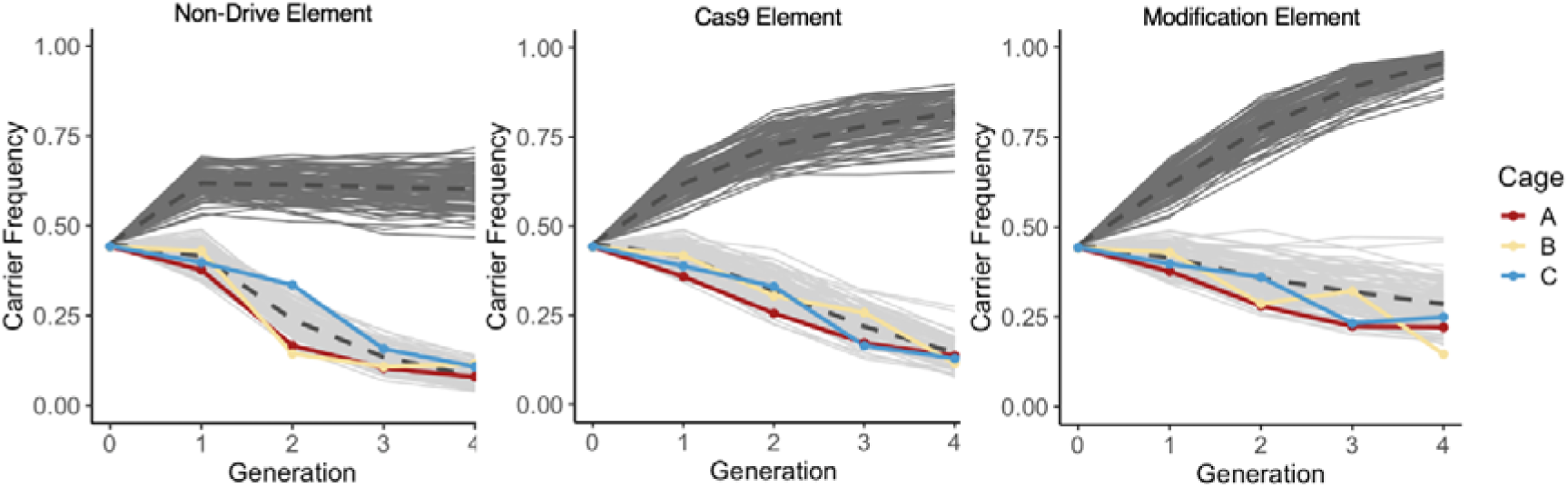
Cage study. Homozygotes for each daisy chain allele were introduced into a cage containing *w^1118^* at an initial frequency of 45%, and the cage was followed for several discrete generations. The carrier frequency (including homozygotes and heterozygotes) for each allele is displayed for each generation. The light grey lines indicate modeling results with drive inheritance parameters from this study and other fitness and resistance parameters inferred from previous studies. The darker grey lines indicate modeling results based on similar parameters but with an additional homozygous fitness multiplier for each daisy allele of 0.211 (heterozygotes received the square root of this value), allowing our model to better match the result of the cage experiment.

To further investigate this unexpected fitness cost, we conducted a fertility test for triple drive heterozygous females. We found that although females carrying all three elements laid the same amount of eggs as *w^1118^*, only 10% of the eggs successfully developed into adults (Fig S4), which was 11.3% of the survival of progeny from *w^1118^* flies (Data Set S4). Actual viability may have been slightly higher, but negatively impacted by poor food vial conditions when many nonviable eggs are laid. Based on our default cage performance parameters, we expected this relative rate to be 27.3% for females, assuming that the female fitness cost entirely affected their fertility (surviving offspring). This takes into account offspring nonviability based on genotype (usually from embryo resistance) in the Cas9 and modification drive target sites. However, if we assume that the fitness effect of our system is entirely in females and thus that the fitness cost in females is doubled compared to our cage expectation (no more negative fitness effects in males), then the relative egg viability compared to *w^1118^* would be 7.4%. Thus, our measured fitness cost in females from reduced egg viability of 11.3% is consistent with expectations based on our cage experiment of 7.4% (all fitness costs in females) and 27.4% (fitness costs divided evenly between females and males), supporting the notion that the fitness reduction was not due to an unknown factor that only affected our cage study and not our egg viability experiment.

## Discussion

In our study, we modeled the performance of daisy chain gene drives for population suppression. We found that they could perform well with even a single (albeit large) release in panmictic populations, but their self-limiting nature prevented success in spatial environments outside the immediate release area. We also demonstrated modification and suppression daisy chains experimentally in fruit flies. Inheritance could be biased successfully, but unexpected fitness costs prevented the system from increasing in frequency in mixed populations.

Previous modeling of daisy chain gene drives focused mostly on modification drives in panmictic populations^44,48^. Consistent with these, our model indicates that reducing the cut rate and fitness will impair the suppression daisy chains significantly, more so than self-sustaining homing drives, and this impact was even greater in suppression systems compared to modification systems. Specifically, the suppression system has thresholds of drive conversion rate and relative release size, and successful population elimination only occurs above these thresholds. In our simulation where the low-density growth rate was set to 6 and a concave density growth curve, the threshold for drive conversion rate was 70%, and the threshold for relative release size was 60%, but these would higher for more robust populations. This highlights the increased confinement level of suppression daisy chains, as limited migration would thus be insufficient to cause species extinction in neighboring populations. This is in contrast to modification daisy drives, which can be highly invasive under some circumstances^49^.

Although a previous study showed that increased maternal deposition/embryo resistance can reduce the invasiveness of daisy chain drives^48^, we found that this had almost no impact on suppression daisy chains. This is likely because suppression daisy drives rely on rapid inducement of population sterility for success (before the drive declines in frequency).

Though embryo resistance prevents future drive conversion, it also increases the frequency of sterile female progeny, even if they have just one drive allele (or even if they lack any). Thus, our result shows that these factors likely cancel out in determining population elimination outcomes. One major challenge for developing gene drive systems in at least some species is controlling Cas9 expression to minimize embryo resistance^14,68^. This insensitivity allows daisy chain designs, particularly suppression drives, to focus on enhancing drive conversion efficiency and minimize fitness costs from somatic expression, thus potentially allowing for easier development in different organisms.

In continuous space, we found that suppression daisy chains cannot significantly spread at all. Unlike panmictic populations in which drive individuals we release are perfectly mixed with wild type populations, we only released transgenics in a limited area in the spatial simulation. In this circumstance, the ability for a gene drive system to spread in continuous space largely depends on whether the frequency of drive alleles can increase at the boundary between the drive and wild-type areas^58,69^. For self-limiting gene drives like daisy chains, their frequency at the boundary is likely to fall rapidly, and for suppression systems, there will be very little migration of alleles from areas of the initial release due to the reduced population in this region. Therefore, suppression daisy chains designs cannot be expected eliminate a large population in a spatial area unless releases take place over nearly the entire region. This is an important contrast to confined frequency-dependent gene drive systems which can sometimes spread outside the release region if they are released at sufficiently high frequency to successfully establish.

In this study, we constructed modification and suppression daisy chain systems in a model organism as a proof-of-principle. Considering that functional resistance alleles can outcompete the drive allele, all the target sites we choose in this study were essential genes targeted by multiple gRNAs, which are usually sufficient to prevent functional resistance^14,34,38^. Each allele also showed adequate inheritance results individually and when all three elements were combined together, demonstrating the potential utility of daisy chain systems despite possible inefficiency from saturation of Cas9 with many gRNAs.

However, our cage experiment results indicate that daisy chain gene drives may face larger obstacles than expected. We expected a moderate fitness cost in our Cas9 element^61^ and little to no fitness cost in our modification element^34^ and non-drive element. Yet, our egg viability experiments indicate that triple heterozygote females have a 90% decrease in egg hatching rates, substantially larger than expected even when considered nonviability due to nonfunctional resistance alleles at two of our daisy element sites. In cage experiments, even the modification element itself did not increase in frequency, and models only matched experimental data when we included much larger fitness costs (that were approximately in-line with our egg viability experiment). It remains unclear exactly what may be causing these fitness costs. It is possible that previous studies underestimated fitness costs, but even using the bottom of our 95% confidence intervals would only explain part of them. Fitness costs may have been increased due to different food type compared to previous studies. The complex structure of a daisy chains, with multiple transgenes, may also result in sufficiently high transgene expression to alone start to have notable effects. Additional fitness costs could come from off-target effects^70^ that played a smaller role in situations with rapid drive spread, simultaneous cleavage at different genomic sites, or other unknown aspects.

In summary, our study used modeling to explore the key parameters that need to be considered when designing a daisy chain gene drive system for population suppression. We successfully engineered two daisy chains that achieved drive conversion in *D. melanogaster*, though they suffered from unknown fitness costs in cage experiments. Our results are nonetheless promising for future construction of self-limiting suppression systems, providing more options that may be even stronger than single-element system designs^47^ by avoiding the need for multiple releases. However, future experiments will need to assess whether multiple drive elements can be used together in some situations without prohibitive fitness costs.

## Supporting information

Supplementary Data Sets

## Acknowledgements

This study was supported by Peking University, the Center for Life Sciences, and the National Natural Science Foundation of China (grants 32270672 and W2432018). The cluster-based data collection was assisted by High-Performance Computing Platform of the Center for Life Science at Peking University.

## Supplemental Information Figure

**Figure S1.**
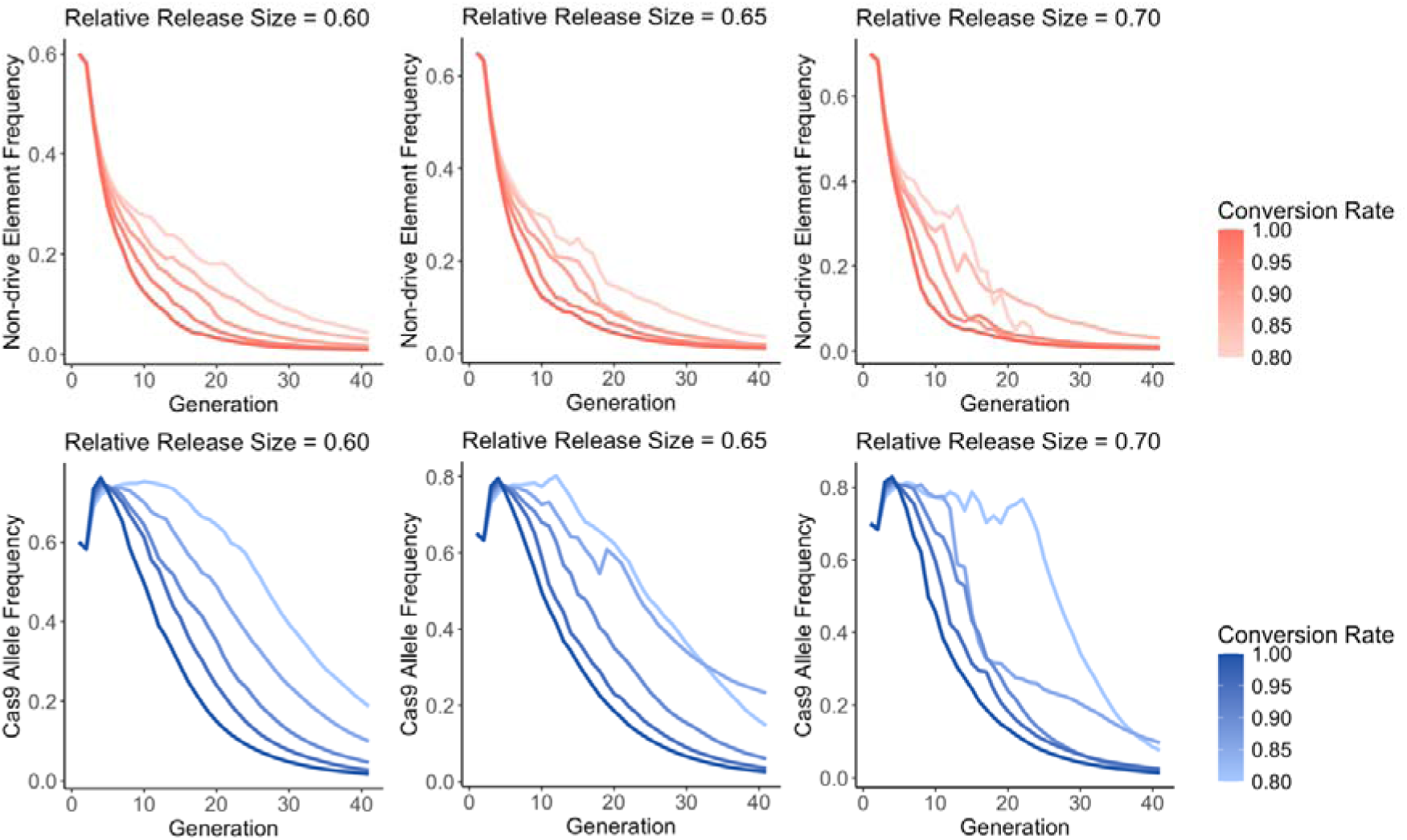
For daisy suppression drives, we measured the **a-c** non-drive element frequency and **d-f** Cas9 element frequency under different drive conversion rates. Each line represents the mean value of 10 replicates.

**Figure S2.**
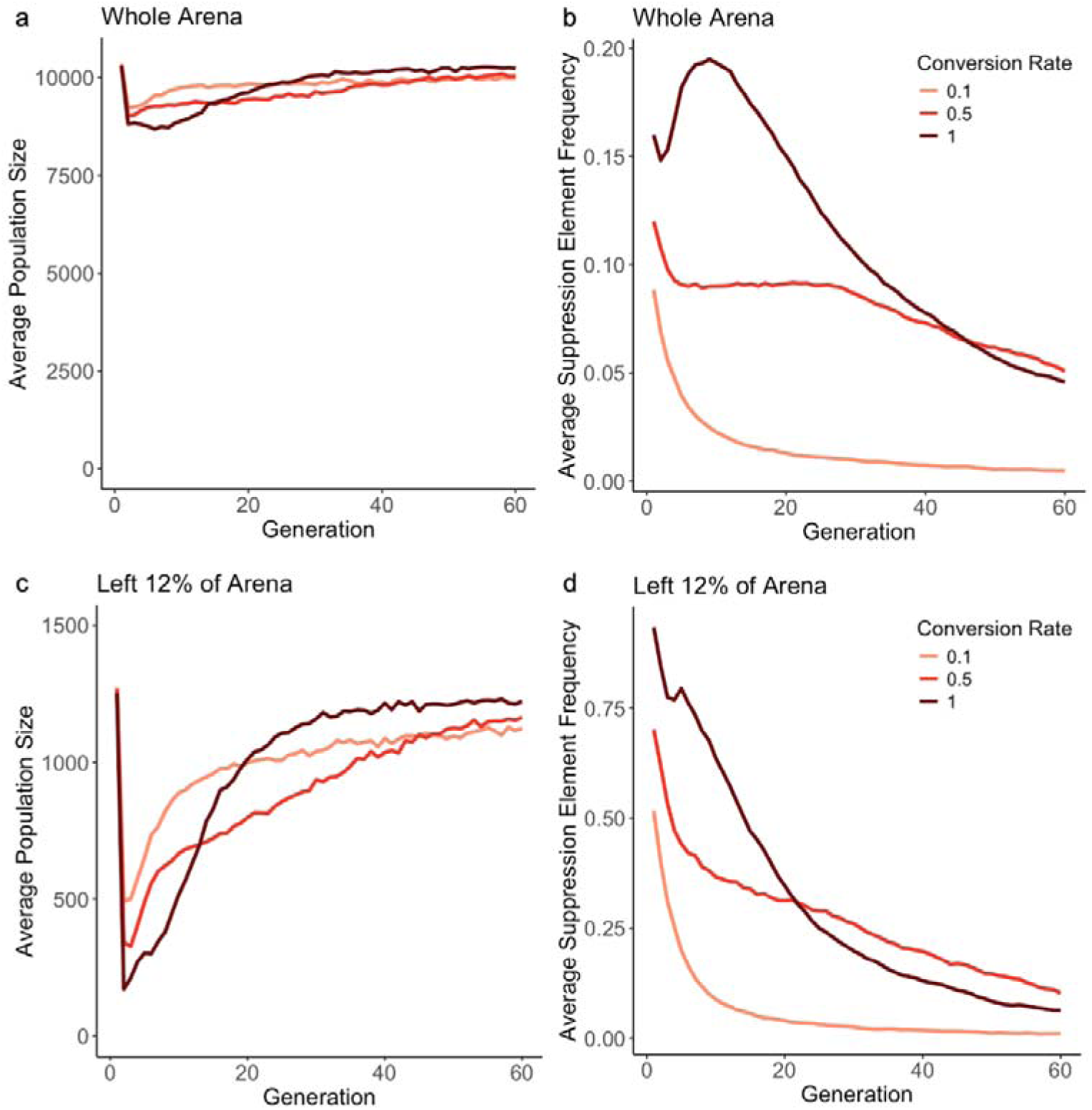
We released a daisy drive in the left 20% of a one-dimensional spatial arena. In the **a-b** left 12% of the arena or **c-d** the whole area, we show the **a,c** average population size and the **b,d** suppression element frequency based on of 20 replicates for each value of drive conversion rate.

**Figure S3.**
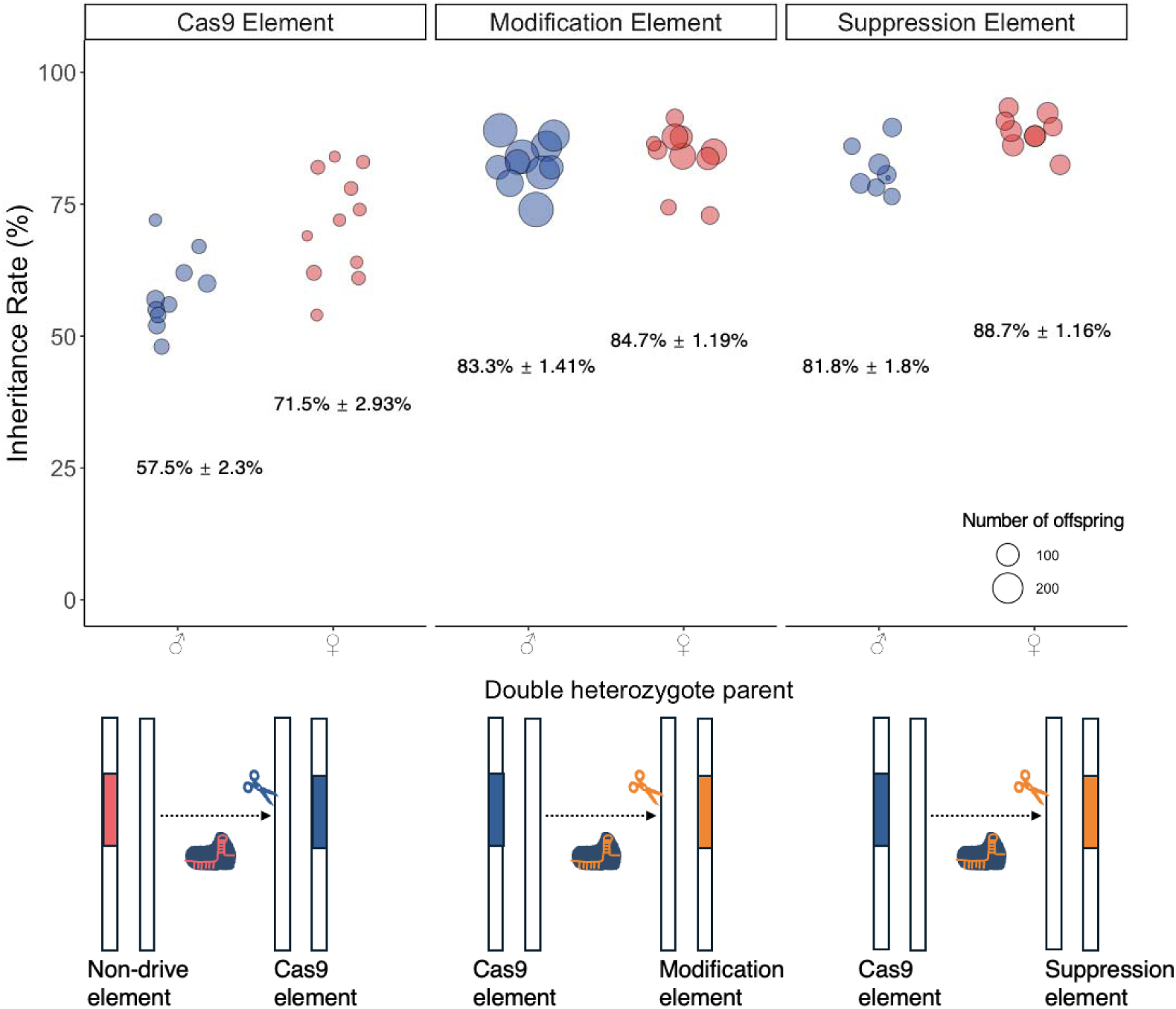
Inheritance rates in double heterozygotes. The inheritance rate of the Cas9 element was evaluated by crossing individuals that were heterozygous for the non-drive element and Cas9 element with *w^1118^* flies. The inheritance rate of the modification and suppression elements was evaluated by crossing individuals that were heterozygous for the Cas9 element and the modification/suppression element with *w^1118^* flies. Each data point represents the inheritance rate observed in one vial containing one drive parent, and the size of each dot represents the clutch size representing the total number of adult offspring phenotype. Also shown are the mean and standard error.

**Figure S4.**
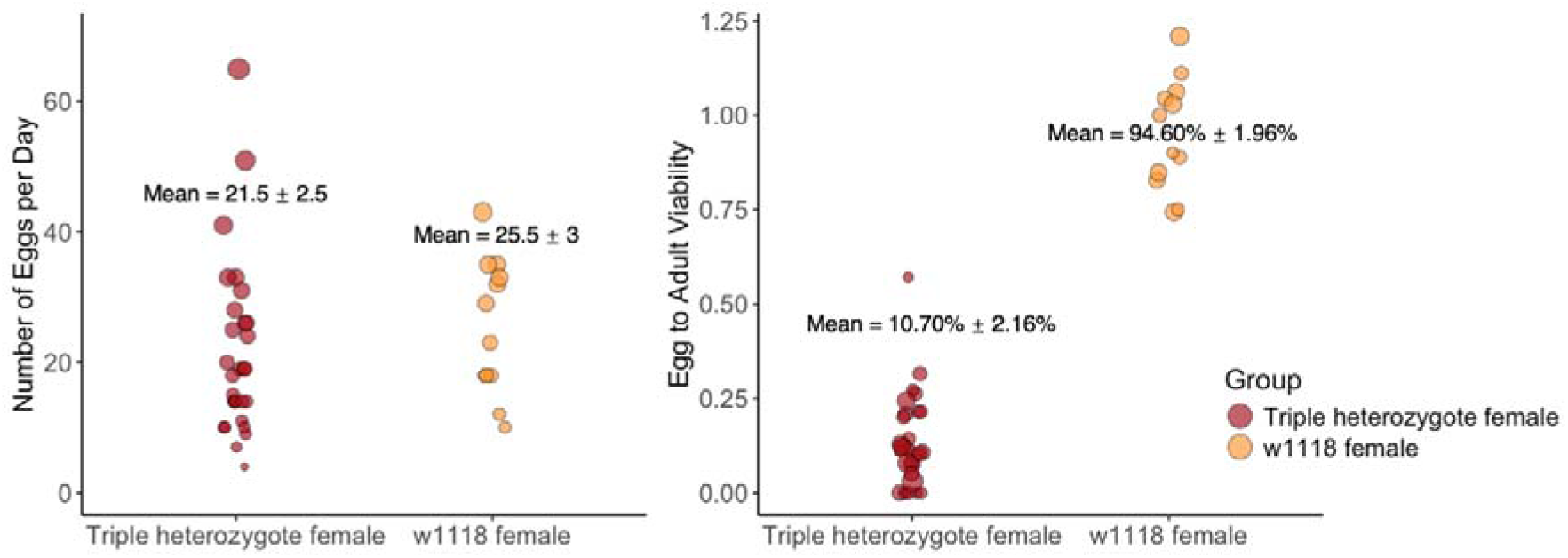
Triple heterozygous females with the modification drive (from crosses between transgenic males and *w^1118^* females) and *w^1118^* females were crossed to *w^1118^* males. We measured the average number of eggs laid per day, and the viability of these eggs (note that undercounting of eggs may result in individual vial viabilities above 1.). Each data point represents an observation in one vial containing one drive parent. The size of each dot represents the total number of eggs laid per day. Also shown are the mean and standard error.

**Table S1.** Cage model parameters.

| Parameters | Values |
| --- | --- |
| Population carrying capacity | 1221 |
| Drive conversion rate (Cas9 element) | 0.49 |
| Drive conversion rate (modification element) | 0.71 |
| Embryo cut rate (Cas9 element) | 0.63 |
| Embryo cut rate (modification element) | 0.29 |
| Fitness (non-drive element) | 0.95 |
| Fitness (Cas9 element) | 0.853 |
| Fitness (modification element) | 0.95 |
| Germline resistance rate (Cas9 element) | 0.398 |
| Germline resistance rate (Modification element) | 0.0934 |
| Functional resistance (Cas9 element) | 0.33 |
| Functional resistance (Modification element) | 0 |

